# The genetic regulation of thyroid hormones on immune and hematopoietic homeostasis: an integrative Mendelian randomization and colocalization study

**DOI:** 10.64898/2026.11.16.767148

**Authors:** Emmanuel Owusu Ansah, Kwadwo Fosu, Isaac Kofi Adu, Aaron Lanidune, Emmanuel Addai Gyabaah, Dorcas Antwi, Philip Boakye Bonsu, Foster Kyei

## Abstract

**Background:** Hematological traits are closely linked to immune and inflammatory processes implicated in thyroid dysfunction. Although observational studies have reported associations between blood-cell traits and thyroid disorders, whether these relationships reflect causal mechanisms remains unclear.

**Methods:** We performed a two-sample Mendelian randomization (MR) analysis to investigate the causal effects of 25 genetically predicted hematological traits on hypothyroidism and hyperthyroidism using genome-wide association study (GWAS) summary statistics from the UK Biobank. Independent single-nucleotide polymorphisms (SNPs) were selected as instrumental variables and harmonized before analysis. Causal estimates were primarily obtained using the inverse-variance weighted (IVW) method and complemented by MR-Egger, weighted median, weighted mode, simple mode, MR-PRESSO, and MR-robust adjusted profile score (MR-RAPS) analyses. Sensitivity analyses assessed heterogeneity, horizontal pleiotropy, reverse causality, and directionality. Significant findings were replicated using independent FinnGen summary statistics. Multivariable MR and Bayesian colocalization analyses were subsequently performed to investigate significant associations further.

**Results:** Among the 25 hematological traits evaluated, eosinophil-related traits showed the strongest associations with hypothyroidism. Genetically predicted eosinophil counts were associated with an increased risk of hypothyroidism (odds ratio [OR] = 1.25, 95% confidence interval [CI]: 1.11–1.42, *P* = 4.0 × 10 , FDR = 0.002), while combined eosinophil–basophil counts showed similar associations (OR = 1.24, 95% CI: 1.10–1.40, *P* = 5.0 × 10 , FDR = 0.002). These findings were consistently replicated in FinnGen and remained significant after adjustment for basophil traits, body mass index, and C-reactive protein in multivariable MR analyses. No significant associations were observed for hyperthyroidism. Bayesian colocalization identified strong evidence of a shared genetic signal at the rs653178 locus on chromosome 12 (PP.H4 = 0.994).

**Conclusions:** This study provides genetic evidence supporting a causal role for eosinophil-related traits in hypothyroidism susceptibility. The consistency of the findings across complementary Mendelian randomization approaches, independent replication, multivariable analyses, and colocalization strengthens the evidence that eosinophil-related immune mechanisms contribute to hypothyroidism susceptibility.

## 1 Introduction

Hypothyroidism and hyperthyroidism are among the most common endocrine disorders worldwide and contribute substantially to morbidity through their effects on cardiovascular, metabolic, neurological, and immune function [**1**]. Hypothyroidism affects approximately 3–5% of the general population, whereas hyperthyroidism affects about 1–2%, with autoimmune thyroid diseases, particularly Hashimoto’s thyroiditis and Graves’ disease, accounting for most cases [**2**]. Early identification of biological pathways contributing to thyroid dysfunction is important for improving disease prediction and understanding pathogenesis.

Hematological abnormalities are frequently observed in individuals with thyroid dysfunction. Previous clinical and observational studies have reported alterations in erythrocyte indices, leukocyte populations, platelet parameters, and inflammatory blood-cell traits in patients with hypothyroidism and hyperthyroidism [**3–5**]. Thyroid hormones regulate hematopoiesis through direct effects on hematopoietic progenitor cells and indirectly through erythropoietin signaling and immune modulation [**6,7**]. Beyond these endocrine effects, autoimmune thyroid diseases are characterized by chronic immune activation, suggesting that circulating hematological traits may reflect broader immune and inflammatory processes involved in disease development [**8**].

Among hematological traits, immune-cell populations such as eosinophils and basophils have attracted increasing attention because of their roles in cytokine signaling, inflammatory regulation, and tissue homeostasis rather than simply allergic responses [**9,10**]. However, whether alterations in hematological traits contribute to thyroid dysfunction or merely represent secondary consequences of disease remains unclear. Most evidence linking hematological traits with thyroid dysfunction has been derived from observational and case-control studies, which are susceptible to residual confounding, reverse causation, medication effects, and disease-related changes in blood-cell composition [**11**]. Because thyroid dysfunction itself influences hematopoiesis and immune-cell distribution, conventional epidemiological studies cannot reliably distinguish causal relationships from downstream physiological consequences.

Mendelian randomization (MR) provides an alternative approach for evaluating potential causal relationships by using genetic variants associated with exposures as instrumental variables, thereby reducing confounding and reverse causation [**12**]. The availability of large-scale genome-wide association studies (GWAS) for hematological traits and thyroid disorders now enables systematic evaluation of these relationships at the genetic level. Furthermore, complementary analyses, including multivariable MR and Bayesian colocalization, can help determine whether observed associations are independent of correlated traits and whether exposure and outcome share common genetic signals.

In the present study, we applied a comprehensive two-sample MR framework to investigate the causal effects of 25 genetically predicted hematological traits on hypothyroidism and hyperthyroidism. Significant associations were further evaluated using independent replication analyses, multivariable MR, and Bayesian colocalization. We hypothesized that specific immune-related hematological traits would contribute to thyroid dysfunction susceptibility and that integrating complementary genetic approaches would strengthen causal inference.

## 2 Method

### 2.1 Study Design

We investigated the causal effects of thyroid hormones on hematological traits and sought to characterize the genetic architecture underlying the identified associations. An integrative analytical framework was applied using genome-wide association study (GWAS) summary statistics from individuals of European ancestry. Two-sample Mendelian randomization (MR) was first used to evaluate the effects of thyroid-stimulating hormone (TSH), free thyroxine (FT4), and free triiodothyronine (FT3) on 25 hematological traits. Significant associations were subsequently examined using sensitivity analyses, reverse MR, and multivariable MR. An overview of the MR study design is presented in **Fig. 1**.

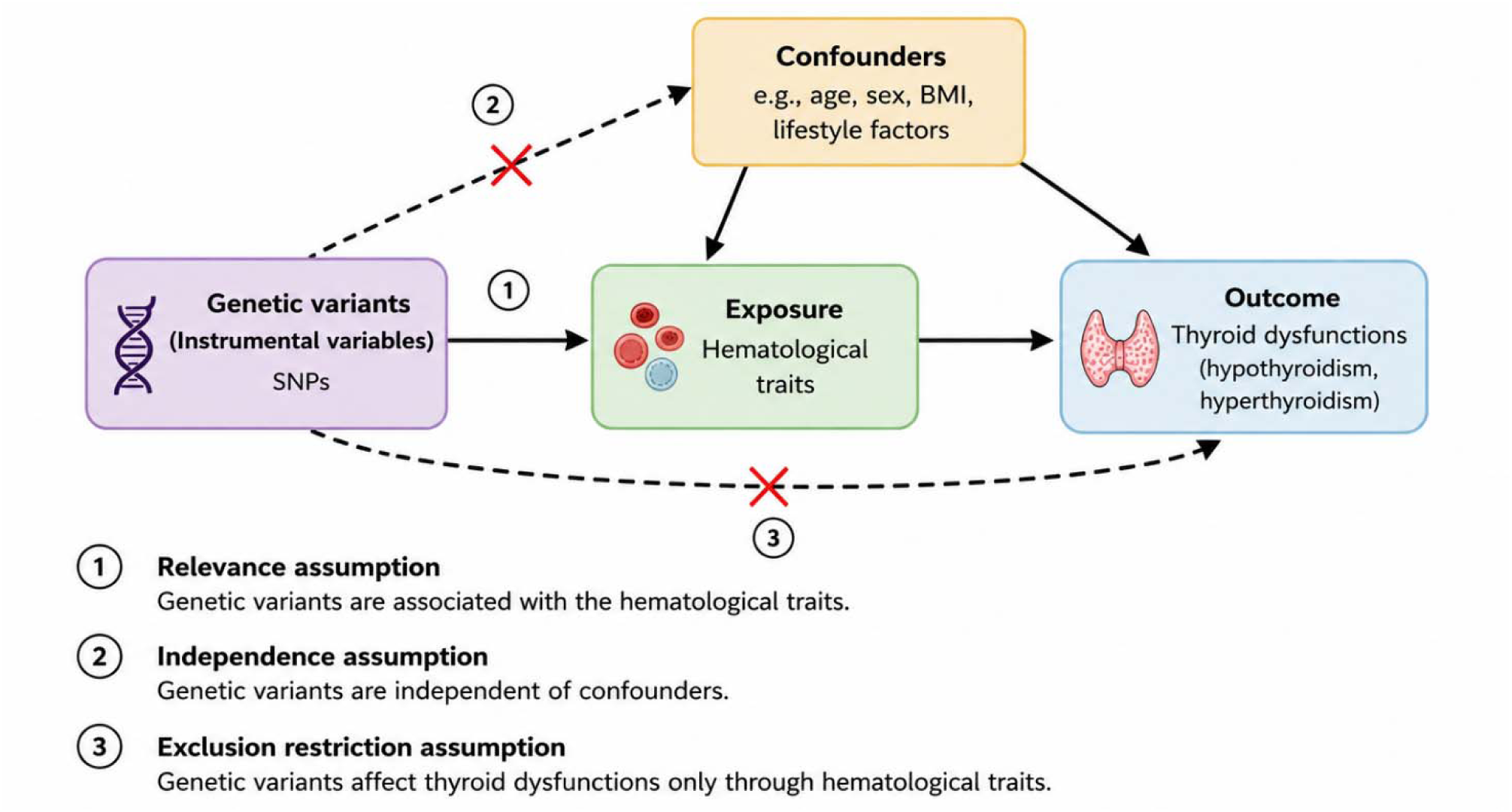

### 2.1 GWAS data for hematological traits

Genetic instruments for hematological traits were obtained from large-scale GWAS of blood-cell indices conducted in 173,480 individuals of European ancestry, as reported by Astle et al. and accessed through the IEU OpenGWAS Project **[15]**. The study included participants from the UK Biobank, UK BiLEVE, and INTERVAL cohorts. A total of 25 hematological traits representing red blood cell, white blood cell, and platelet indices were included in the present analysis. Detailed information on participant recruitment, phenotype measurement, genotyping, imputation, and quality-control procedures has been described previously **[15]**.

#### 2.2.2 GWAS Data for Thyroid Hormones

Genetic association data for thyroid-stimulating hormone (TSH), free thyroxine (FT4), and free triiodothyronine (FT3) were obtained from the ThyroidOmics Consortium genome-wide association study (GWAS) of individuals of European ancestry reported by Sterenborg et al. [16]. Summary statistics included up to 247,107 participants for TSH, 189,425 for FT4, and 173,925 for FT3. These GWAS meta-analyses combined data from multiple population-based cohorts and included rigorous quality-control, genotype imputation, and association analyses as described in the original publication [16]. All analyses in the present study were restricted to individuals of European ancestry to minimize bias due to population stratification.

### 2.3 Instrumental Variable Selection

Genetic instruments for each hematological trait were selected from the corresponding GWAS summary statistics. Single-nucleotide polymorphisms (SNPs) associated with each exposure at genome-wide significance (P < 5 × 10 ) were retained. To ensure independence between instruments, linkage disequilibrium (LD) clumping was performed using the European reference panel from the 1000 Genomes Project, applying a threshold of r² < 0.001 within a 10,000 kb window. Instrument strength was assessed using F-statistics, and only instruments with F > 10 were retained [**18**]. Exposure and outcome datasets were harmonized to ensure consistent alignment of effect alleles. Palindromic SNPs with ambiguous strand orientation were excluded, and proxy variants in high LD (r² > 0.8) were used when necessary.

### 2.4 Mendelian Randomization Analysis

Two-sample MR analyses were performed to estimate the effects of hematological traits on hypothyroidism and hyperthyroidism. The inverse variance weighted (IVW) method was used as the primary analytical approach. Sensitivity analyses were conducted using MR-Egger regression, weighted median, weighted mode, simple mode, MR-RAPS, and MR-PRESSO **[14, 19]**. Heterogeneity among genetic instruments was assessed using Cochran’s Q statistic. Horizontal pleiotropy was evaluated using the MR-Egger intercept test, while outlier variants were identified using MR-PRESSO. Leave-one-out analyses were performed to assess the influence of individual SNPs on the overall causal estimates **[20, 21]**. Replication analyses were conducted using independent GWAS summary statistics from FinnGen [**22**]. Directionality was assessed using the Steiger test, and reverse MR analyses were performed to evaluate the possibility of reverse causation. Effect estimates were reported as odds ratios (ORs) or beta coefficients with corresponding 95% confidence intervals (CIs). This study adhered to the STROBE-MR checklist for transparent reporting of MR analyses [**23**].

### 2.5 Multivariable Mendelian Randomization

Multivariable Mendelian randomization (MVMR) analyses were performed to evaluate whether the associations identified in the primary MR analyses were independent of related hematological, inflammatory, and metabolic traits [**24**]. In the primary model, eosinophil counts and basophil-related traits were analyzed jointly. In a secondary model, eosinophil counts were evaluated together with C-reactive protein (CRP) and body mass index (BMI). Genetic instruments associated with at least one exposure were combined and pruned for independence using an LD threshold of r² < 0.001. Following harmonization, direct effects were estimated using the multivariable IVW method. Conditional F-statistics were calculated to assess instrument strength.

### 2.6 Colocalization Analyses

Bayesian colocalization analyses were performed to determine whether associations between hematological traits and thyroid dysfunction were driven by shared genetic variants [**25**]. Analyses were conducted at genomic regions showing overlapping association signals between exposure and outcome datasets. Variants located within ±500 kb of lead SNPs were extracted and harmonized across datasets. Colocalization analyses were performed using default prior probabilities (p = 1 × 10 , p = 1 × 10 , and p = 1 × 10 ). Hypothyroidism was modeled as a case–control trait with a population prevalence of 0.1. Evidence for colocalization was assessed using posterior probabilities, with PP.H4 ≥ 0.75 considered indicative of a shared causal variant.

## Results

### 3.1 Instrument Selection and Harmonization

After linkage disequilibrium clumping, 49, 5, and 6 independent genetic variants were selected as instrumental variables for TSH, FT4, and FT3, respectively (**Supplementary Table S1**). The selected instruments exhibited strong instrument strength, with overall F-statistics well above the conventional threshold of 10, thereby minimizing the likelihood of weak-instrument bias. Harmonization of exposure and outcome summary statistics ensured consistent alignment of effect alleles across all analyses. The final harmonized datasets were subsequently used to estimate the causal effects of thyroid hormone traits on hematological parameters using two-sample Mendelian randomization.

### 3.2 Associations Between Thyroid Hormones and Hematological Traits

Primary Mendelian randomization analyses were performed to investigate the causal effects of genetically predicted thyroid-stimulating hormone (TSH), free thyroxine (FT4), and free triiodothyronine (FT3) on 25 hematological traits (Figure 2; Supplementary Table S2). Overall, FT3 demonstrated the strongest and most consistent associations with hematological indices, whereas TSH and FT4 showed comparatively limited evidence of association. For TSH, only red blood cell count showed a nominal inverse association (OR = 0.933, 95% CI: 0.873–0.997, *P* = 0.040), while the remaining hematological traits were not significantly associated after correction for multiple testing. Similarly, FT4 showed nominal positive associations with eosinophil count (OR = 1.365, 95% CI: 1.119–1.664, *P* = 0.002), eosinophil–basophil count (OR = 1.336, 95% CI: 1.108–1.611, *P* = 0.002), eosinophil percentage of white cells (OR = 1.312, 95% CI: 1.071– 1.608, *P* = 0.009), and an inverse association with immature reticulocyte fraction (OR = 0.803, 95% CI: 0.676–0.954, *P* = 0.012). However, none of these associations remained statistically significant after Bonferroni correction.

**Figure 2.**
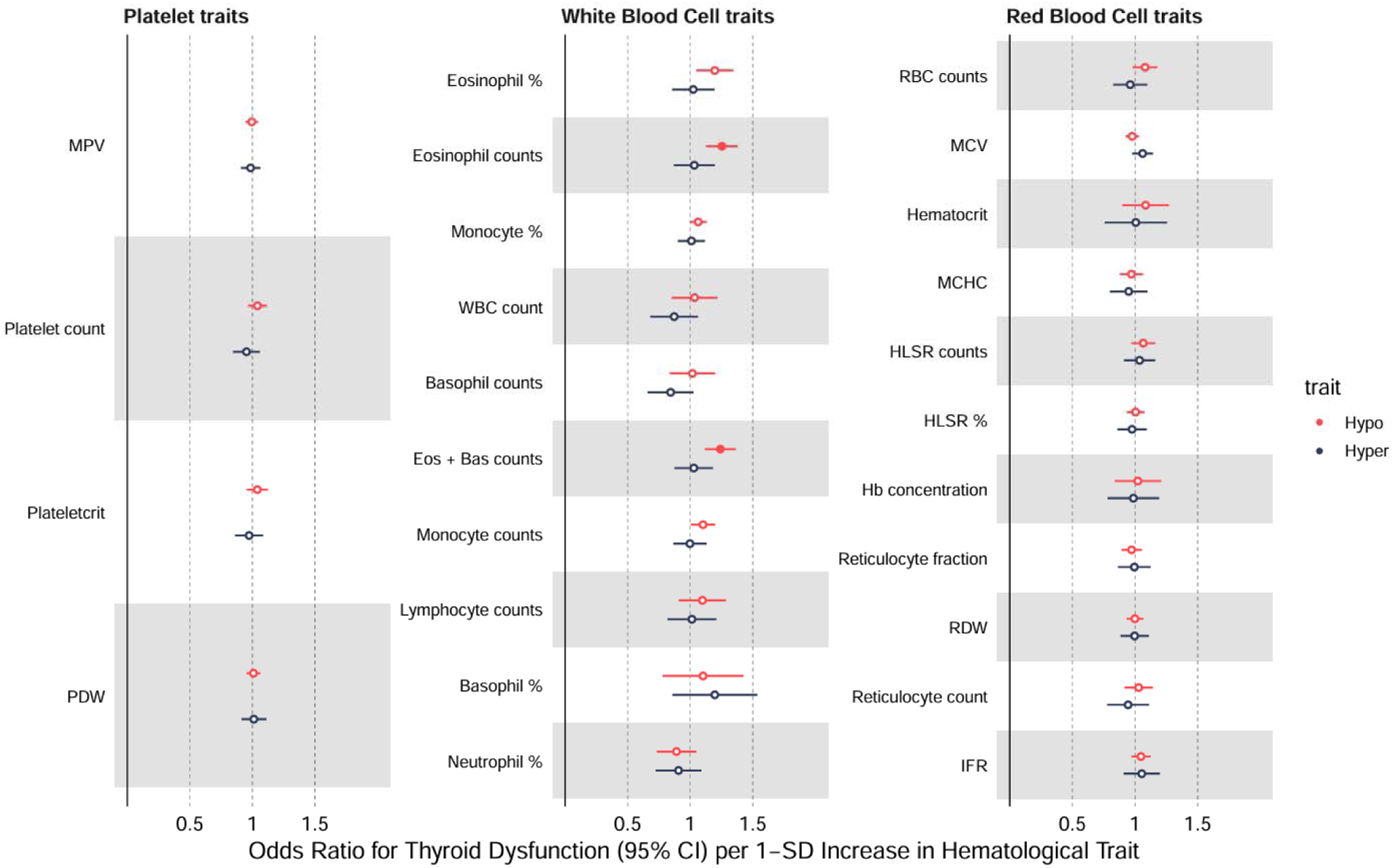
Mendelian randomization estimates for the associations between hematological traits and thyroid dysfunction. Odds ratios (ORs) and 95% confidence intervals (CIs) are shown for inverse-variance weighted analyses.

In contrast, FT3 exhibited the greatest number of significant associations. Eight hematological traits were associated with genetically predicted FT3 at the nominal significance level (*P* < 0.05), including hemoglobin concentration, hematocrit, mean corpuscular volume, red blood cell count, platelet count, plateletcrit, mean platelet volume, and basophil percentage of white cells. Of these, five associations remained significant after Bonferroni correction (Table 1). Higher genetically predicted FT3 levels were associated with increased red blood cell count (OR = 1.178, 95% CI: 1.095–1.267, *P* = 1.05 × 10 ) and hemoglobin concentration (OR = 1.136, 95% CI: 1.055–1.222, *P* = 6.62 × 10 ), whereas inverse associations were observed for mean corpuscular volume (OR = 0.889, 95% CI: 0.846–0.935, *P* = 3.62 × 10 ), platelet count (OR = 0.906, 95% CI: 0.861–0.954, *P* = 1.60 × 10 ), and basophil percentage of white cells (OR = 0.915, 95% CI: 0.872–0.961, *P* = 3.89 × 10 ).

**Table 1.** Primary Mendelian randomization estimates for significant hematological traits associated with free triiodothyronine (FT3). Odds ratios (ORs), 95% confidence intervals (CIs), and P-values were estimated using the inverse-variance weighted (IVW) method. Associations were evaluated across 25 hematological traits. OR, odds ratio; CI, confidence interval; IVW, inverse-variance weighted.

| Hematological trait | MR method | OR | 95% CI | P-value |
|---|---|---|---|---|
| Red blood cell count | IVW | 1.184 | 1.109–1.265 | 4.83 × 10□□ |
| Mean corpuscular volume | IVW | 0.895 | 0.848–0.944 | 4.21 × 10□□ |
| Hemoglobin concentration | IVW | 1.152 | 1.065–1.247 | 3.98 × 10□□ |
| Platelet count | IVW | 0.912 | 0.863–0.963 | 1.00 × 10□³ |
| Basophil percentage of white cells | IVW | 0.926 | 0.879–0.976 | 4.15 × 10□³ |

### 3.3 Sensitivity Analyses Support the Robustness of the FT3 Associations

Sensitivity analyses supported the robustness of the five Bonferroni-significant associations between genetically predicted FT3 and hematological traits. Weighted median and MR-RAPS analyses produced effect estimates that were highly consistent with the primary inverse-variance weighted (IVW) results, while MR-Egger regression, simple mode, and weighted mode generally yielded estimates in the same direction, albeit with wider confidence intervals and reduced statistical significance (**Table 2; Supplementary Table S3**). MR-Egger intercept tests showed no evidence of directional horizontal pleiotropy for any of the five significant associations (all *P* > 0.05). Cochran’s Q test indicated no significant heterogeneity for mean corpuscular volume, platelet count, or basophil percentage of white cells. In contrast, modest heterogeneity was observed for red blood cell count and hemoglobin concentration (**Supplementary Table S3**). MR-PRESSO analyses detected no influential outlier variants (**Supplementary Table S4**). Steiger directionality analyses consistently supported the inferred causal direction from FT3 to the hematological traits, with the instrumental variants explaining substantially more variance in FT3 than in the corresponding hematological outcomes (**Supplementary Table S5**). Reverse Mendelian randomization analyses provided limited evidence for reverse causality (**Supplementary Table S6**).

**Table 2.**
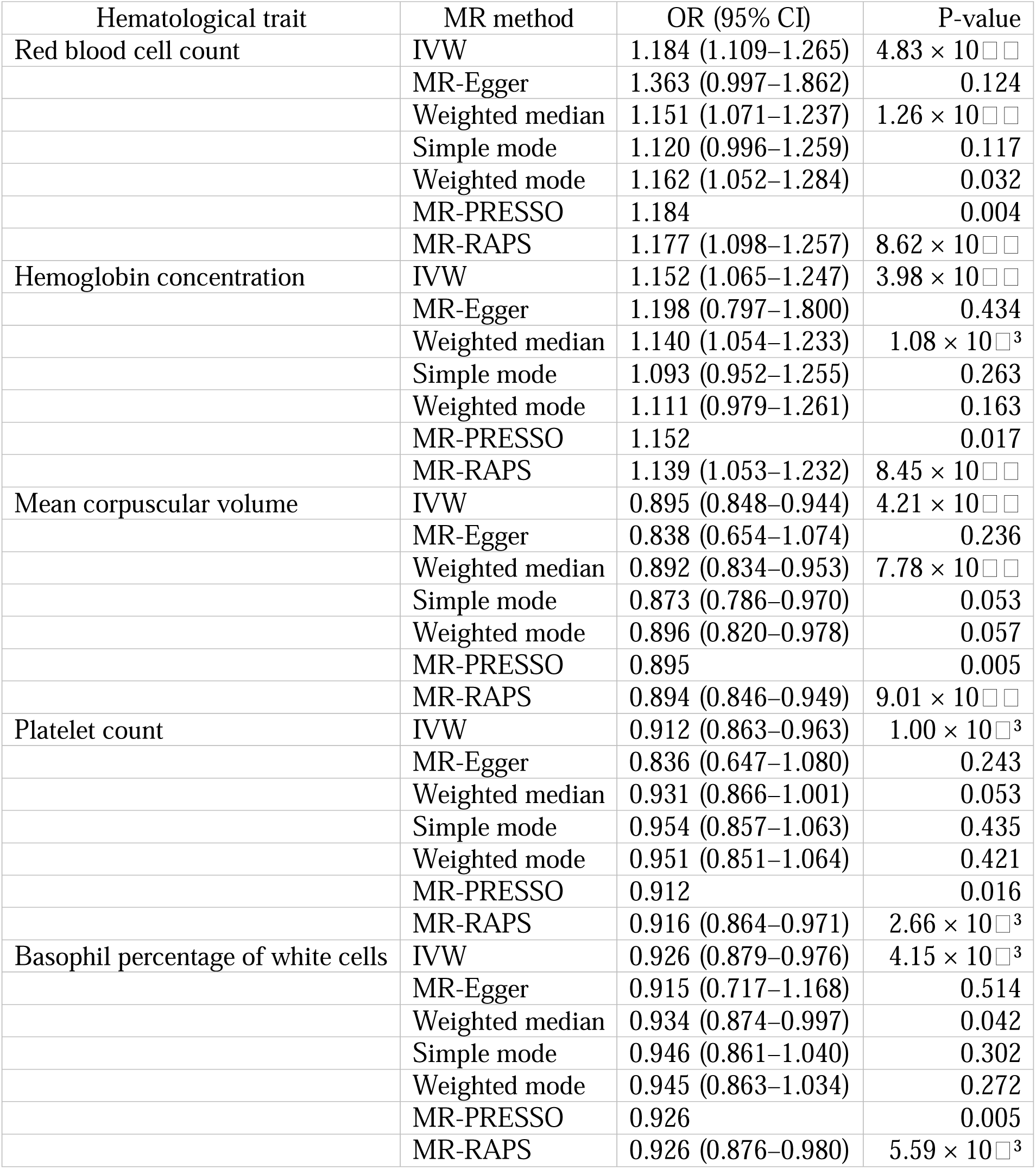
Sensitivity analyses of the associations between genetically predicted free triiodothyronine (FT3) levels and significant hematological traits using complementary Mendelian randomization methods.

| Hematological trait | MR method | OR (95% CI) | P-value |
|---|---|---|---|
| Red blood cell count | IVW | 1.184 (1.109–1.265) | 4.83 × 10□□ |
|  | MR-Egger | 1.363 (0.997–1.862) | 0.124 |
|  | Weighted median | 1.151 (1.071–1.237) | 1.26 × 10□□ |
|  | Simple mode | 1.120 (0.996–1.259) | 0.117 |
|  | Weighted mode | 1.162 (1.052–1.284) | 0.032 |
|  | MR-PRESSO | 1.184 | 0.004 |
|  | MR-RAPS | 1.177 (1.098–1.257) | 8.62 × 10□□ |
| Hemoglobin concentration | IVW | 1.152 (1.065–1.247) | 3.98 × 10□□ |
|  | MR-Egger | 1.198 (0.797–1.800) | 0.434 |
|  | Weighted median | 1.140 (1.054–1.233) | 1.08 × 10□³ |
|  | Simple mode | 1.093 (0.952–1.255) | 0.263 |
|  | Weighted mode | 1.111 (0.979–1.261) | 0.163 |
|  | MR-PRESSO | 1.152 | 0.017 |
|  | MR-RAPS | 1.139 (1.053–1.232) | 8.45 × 10□□ |
| Mean corpuscular volume | IVW | 0.895 (0.848–0.944) | 4.21 × 10□□ |
|  | MR-Egger | 0.838 (0.654–1.074) | 0.236 |
|  | Weighted median | 0.892 (0.834–0.953) | 7.78 × 10□□ |
|  | Simple mode | 0.873 (0.786–0.970) | 0.053 |
|  | Weighted mode | 0.896 (0.820–0.978) | 0.057 |
|  | MR-PRESSO | 0.895 | 0.005 |
|  | MR-RAPS | 0.894 (0.846–0.949) | 9.01 × 10□□ |
| Platelet count | IVW | 0.912 (0.863–0.963) | 1.00 × 10□³ |
|  | MR-Egger | 0.836 (0.647–1.080) | 0.243 |
|  | Weighted median | 0.931 (0.866–1.001) | 0.053 |
|  | Simple mode | 0.954 (0.857–1.063) | 0.435 |
|  | Weighted mode | 0.951 (0.851–1.064) | 0.421 |
|  | MR-PRESSO | 0.912 | 0.016 |
|  | MR-RAPS | 0.916 (0.864–0.971) | 2.66 × 10□³ |
| Basophil percentage of white cells | IVW | 0.926 (0.879–0.976) | 4.15 × 10□³ |
|  | MR-Egger | 0.915 (0.717–1.168) | 0.514 |
|  | Weighted median | 0.934 (0.874–0.997) | 0.042 |
|  | Simple mode | 0.946 (0.861–1.040) | 0.302 |
|  | Weighted mode | 0.945 (0.863–1.034) | 0.272 |
|  | MR-PRESSO | 0.926 | 0.005 |
|  | MR-RAPS | 0.926 (0.876–0.980) | 5.59 × 10□³ |

### 3.3 Replication Analyses Confirm Associations Between Eosinophil-Related Traits and Hypothyroidism

Because eosinophil counts and combined eosinophil–basophil counts showed the strongest and most consistent evidence of association with hypothyroidism, subsequent analyses focused on eosinophil-related traits. Replication MR analyses using independent FinnGen summary statistics confirmed the associations observed in the UK Biobank dataset. Genetically predicted eosinophil percentage of white cells (OR = 1.20, 95% CI: 1.06–1.37, P = 0.006), eosinophil counts (OR = 1.23, 95% CI: 1.10–1.38, P < 0.001), and combined eosinophil–basophil counts (OR = 1.24, 95% CI: 1.11–1.38, P < 0.001) were all associated with an increased risk of hypothyroidism (**Figure 3; Supplementary Table S7**).

**Figure 3.**
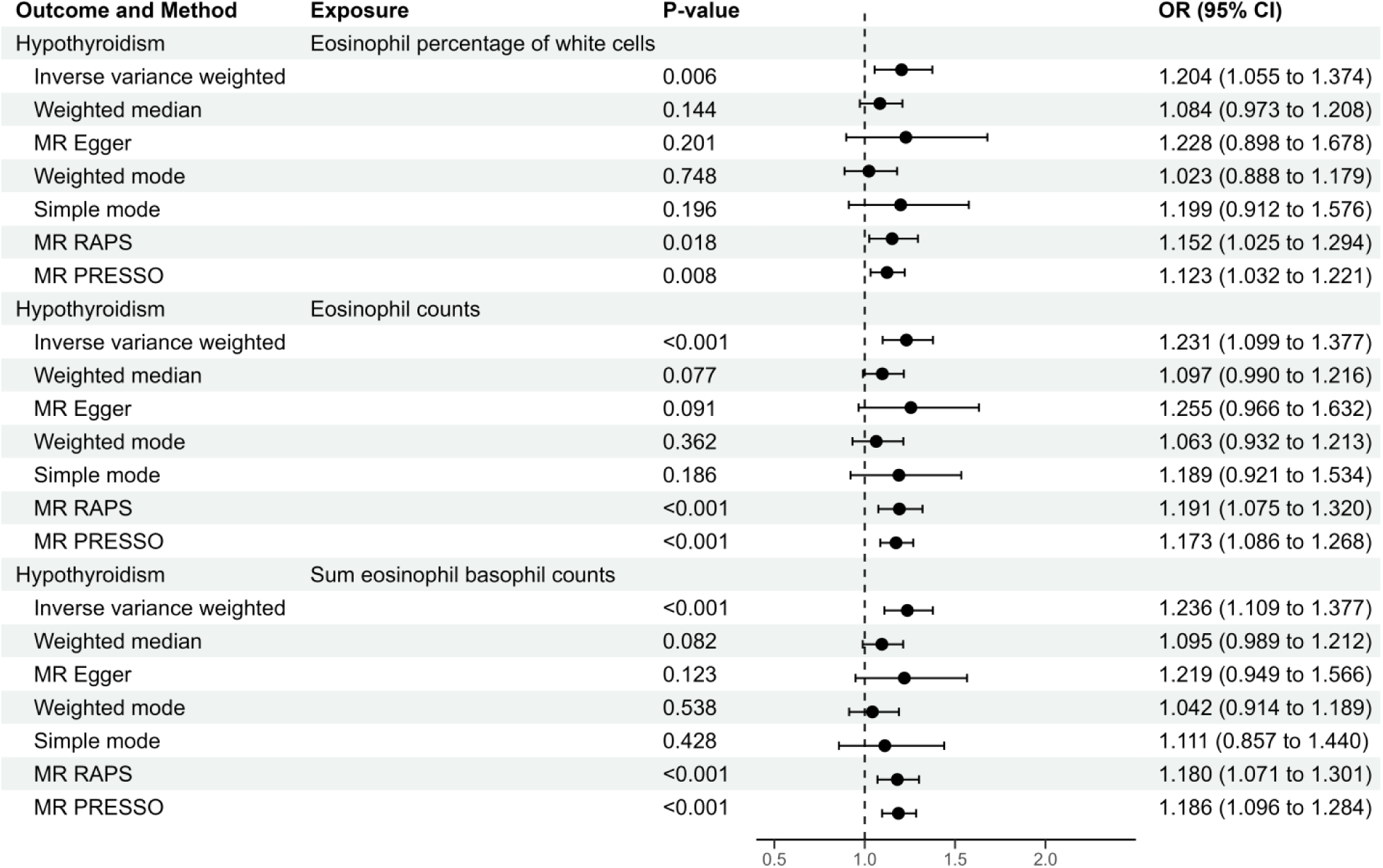
Replication analyses of eosinophil-related traits and hypothyroidism in the FinnGen cohort. Odds ratios (ORs) and 95% confidence intervals (CIs) are shown for inverse-variance weighted analyses.

Sensitivity analyses supported the replication findings. Effect estimates from MR-Egger, weighted median, MR-PRESSO, and MR-RAPS analyses were generally concordant with th primary IVW results (**Figure 3**). Associations for eosinophil counts and combined eosinophil– basophil counts remained significant in MR-RAPS and MR-PRESSO analyses. MR-Egger intercept tests showed no evidence of directional pleiotropy, although Cochran’s Q statistics indicated heterogeneity among instrumental variables (**Supplementary Table S7**). Scatter plots, funnel plots, and leave-one-out analyses showed no evidence that the observed associations were driven by individual variants or directional bias (**Supplementary Figures S2A–S2F**).

### 3.4 Multivariable Mendelian Randomization Supports an Independent Association Between Eosinophil Counts and Hypothyroidism

We then evaluated whether the eosinophil–hypothyroidism association was independent of related immune and inflammatory traits. In a multivariable MR model including eosinophil counts and basophil percentage, eosinophil counts remained significantly associated with hypothyroidism (β = 0.229, SE = 0.078, P = 0.0036), whereas basophil percentage showed no evidence of an independent association (β = −0.043, SE = 0.142, P = 0.764) (**Figure 4; Supplementary Table S8**). To further assess the independence of this relationship, we fitted a second model including eosinophil counts, body mass index (BMI), and C-reactive protein (CRP). The association between eosinophil counts and hypothyroidism remained significant after adjustment for both factors (β = 0.300, SE = 0.052, P = 1.27 × 10 ). CRP was also independently associated with hypothyroidism (β = 0.480, SE = 0.069, P = 1.58 × 10 ¹¹), whereas BMI showed no evidence of association (β = −0.034, SE = 0.043, P = 0.43). Instrument strength was adequate across all multivariable models (**Supplementary Table S8**).

**Figure 4.**
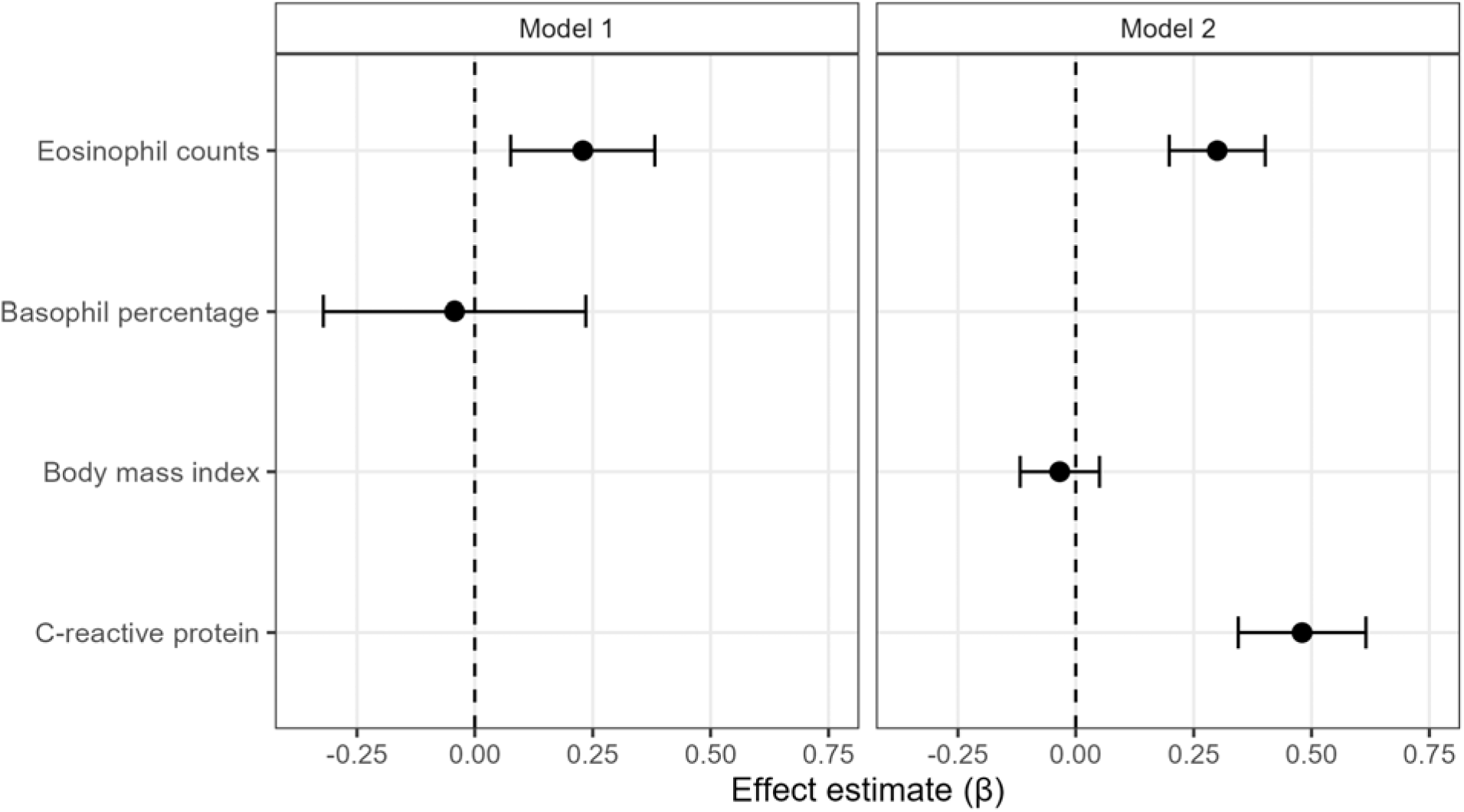
Multivariable Mendelian randomization analyses of eosinophil counts and hypothyroidism. Effect estimates are presented as regression coefficients (β) with corresponding 95% confidence intervals.

### 3.5 Colocalization Analyses Identify a Shared Genetic Signal Between Eosinophil Counts and Hypothyroidism

Given the consistent MR findings, we next examined whether eosinophil counts and hypothyroidism shared genetic association signals. Strong evidence of colocalization was observed at the chromosome 12 rs653178 locus, where a pair of credible sets from eosinophil counts and hypothyroidism showed a high posterior probability of colocalization (PP.H4 = 0.994). Regional association plots demonstrated substantial overlap between eosinophil and hypothyroidism association signals across the locus (**Figure 5**). In contrast, most other loci showed little evidence of colocalization, with posterior probabilities largely supporting distinct association signals for the two traits (**Supplementary Table S9**).

**Figure 5.**
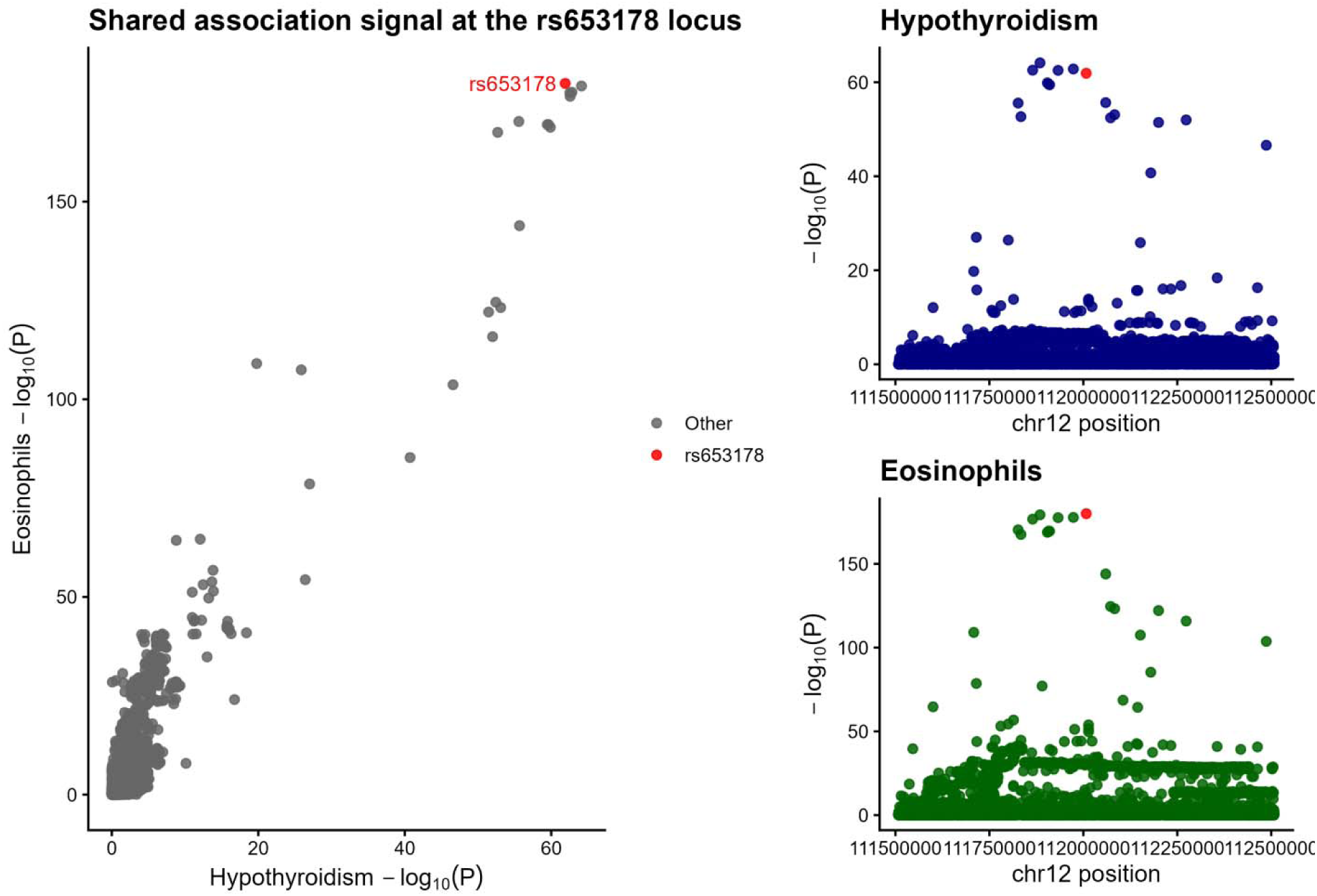
Regional association plots for eosinophil counts and hypothyroidism at the rs653178 locus.

## 4 Discussion

In this study, we investigated the potential causal relationship between genetically predicted hematological traits and thyroid dysfunction using a comprehensive MR framework. Our findings provide genetic evidence that eosinophil-related traits are associated with increased susceptibility to hypothyroidism. Among the 25 hematological traits evaluated, eosinophil counts and combined eosinophil–basophil counts showed the strongest and most consistent associations across discovery, replication, and multivariable analyses, whereas little evidence was observed for the remaining hematological traits. No significant associations were identified for hyperthyroidism, suggesting that the observed effects are specific to hypothyroidism rather than thyroid dysfunction more broadly.

These findings extend previous observational studies reporting hematological abnormalities in patients with thyroid dysfunction, including alterations in leukocyte populations, inflammatory markers, platelet indices, and erythrocyte parameters [**3–5**]. Although such studies consistently demonstrate an association between thyroid disease and hematological traits, they cannot determine whether these abnormalities contribute to disease development or arise as secondary consequences of thyroid dysfunction. Thyroid hormones exert important regulatory effects on hematopoiesis and immune-cell homeostasis, making reverse causation and residual confounding difficult to exclude in conventional epidemiological studies [**2,6**]. By using genetic variants as proxies for lifelong differences in hematological traits, MR substantially reduces these biases and provides stronger evidence that eosinophil-related pathways may contribute to hypothyroidism susceptibility rather than merely accompany established disease.

The observed association between eosinophil-related traits and hypothyroidism is biologically plausible. Although autoimmune thyroid diseases such as Hashimoto’s thyroiditis have traditionally been viewed as disorders driven predominantly by adaptive immunity and autoantibody production, increasing evidence indicates that innate immune mechanisms also contribute substantially to thyroid inflammation and tissue injury [**26**]. Eosinophils are now recognized as multifunctional immune cells involved in cytokine production, inflammatory amplification, oxidative stress regulation, tissue remodeling, and immune-cell communication rather than simply mediators of allergic responses [**27, 28**]. Through the secretion of cytokines, chemokines, lipid mediators, and cytotoxic granule proteins, eosinophils influence the recruitment and activation of macrophages, dendritic cells, mast cells, and lymphocytes, thereby shaping both innate and adaptive immune responses. Persistent eosinophil-associated inflammation may therefore promote chronic immune activation, progressive thyroid tissue injury, and ultimately impaired thyroid hormone production.

The genetic evidence was further strengthened by Bayesian colocalization analyses, which identified a shared association signal at the rs653178 locus on chromosome 12. This region has previously been implicated in immune regulation, hematological traits, and susceptibility to multiple autoimmune diseases [**29, 30**]. The presence of a shared genetic signal supports the hypothesis that common inherited mechanisms contribute to both eosinophil regulation and hypothyroidism susceptibility rather than representing independent associations driven by linkage disequilibrium. Although colocalization alone does not establish biological causality, these findings provide complementary evidence supporting the robustness of the MR results and suggest that immune-regulatory pathways within the chromosome 12q24 region may contribute to the observed relationship.

Additional evidence supporting an independent role for eosinophils was provided by the MVMR analyses. Basophil-related traits, BMI, and CRP were included because they represent biologically plausible factors that could confound or mediate the relationship between eosinophils and hypothyroidism. Basophils share developmental origins and functional characteristics with eosinophils, whereas obesity and systemic inflammation have both been implicated in thyroid dysfunction and immune dysregulation [**31, 32**]. Importantly, eosinophil counts remained significantly associated with hypothyroidism after adjustment for these variables, suggesting that the observed association is unlikely to be explained solely by correlated granulocyte traits, obesity-related pathways, or generalized systemic inflammation. Interestingly, CRP also demonstrated an independent association with hypothyroidism, consistent with previous studies linking chronic low-grade inflammation to autoimmune thyroid disease [**33, 34**]. Together, these findings suggest that multiple immune and inflammatory pathways contribute to hypothyroidism susceptibility, although the persistence of the eosinophil association indicates that its effect is unlikely to be explained by inflammation alone.

The stronger and more consistent associations observed for eosinophil-related traits compared with the other hematological parameters evaluated suggest that specific immune-cell populations may contribute more directly to hypothyroidism susceptibility than broader alterations in hematological physiology. This observation is consistent with increasing recognition that innate immune mechanisms participate alongside adaptive immune responses in the pathogenesis of autoimmune thyroid disease. Future investigations integrating immune-cell phenotyping, single-cell transcriptomics, spatial transcriptomics, and functional experimental models will be valuable for elucidating the molecular mechanisms linking eosinophil biology with thyroid dysfunction and for determining whether eosinophil-associated pathways represent potential targets for disease prediction, prevention, or therapeutic intervention.

This study has several important strengths. First, the MR framework substantially reduces susceptibility to reverse causation and residual confounding that commonly affect observational studies. Second, the use of large GWAS datasets and independent replication analyses in FinnGen increased the robustness and reproducibility of the findings. Third, complementary analytical approaches, including multivariable MR and Bayesian colocalization, provided additional evidence supporting the independence and shared genetic architecture of the observed associations. Finally, the consistency of the findings across multiple MR methods, together with the absence of evidence for substantial directional pleiotropy and the successful replication of the primary associations, strengthens confidence that the observed relationships are unlikely to reflect methodological artifacts.

Several limitations should also be considered. Although extensive sensitivity analyses were performed, residual horizontal pleiotropy cannot be completely excluded. The use of summary-level GWAS data precluded assessment of clinically important factors such as thyroid autoantibody status, disease severity, thyroid hormone concentrations, and treatment history. Furthermore, most participants were of European ancestry, which may limit the generalizability of the findings to other populations. Although Bayesian colocalization identified shared genetic signals, these analyses do not establish the precise biological mechanisms underlying the observed associations. Consequently, experimental validation using cellular and animal models will be necessary to determine how eosinophil-related immune pathways contribute to hypothyroidism development. Future studies in ancestrally diverse populations with richer clinical phenotyping and functional genomic data will be important for confirming these findings and further elucidating the molecular pathways linking hematological traits with thyroid dysfunction.

## Conclusion

This study provides genetic evidence that eosinophil-related traits contribute to susceptibility to hypothyroidism. The consistency of the associations across Mendelian randomization, independent replication, multivariable analyses, and Bayesian colocalization supports a shared genetic basis linking eosinophil biology with hypothyroidism. In contrast, no evidence was observed for a causal relationship between hematological traits and hyperthyroidism, suggesting that the identified associations may be specific to hypothyroidism. These findings highlight the importance of immune-related hematological traits in thyroid disease and demonstrate the value of integrative genetic approaches for strengthening causal inference. Future studies integrating larger multi-ancestry cohorts, detailed immune phenotyping, and functional experimental models are warranted to elucidate the molecular mechanisms linking eosinophil-related pathways to hypothyroidism and to evaluate their potential as biomarkers or therapeutic targets.

## Data Availability Statement

All genome-wide association study (GWAS) summary statistics analyzed in this study are publicly available through the IEU OpenGWAS Project and the FinnGen consortium. Detailed information, including the original studies and accession identifiers for all datasets used, is provided in **Supplementary Table S1**. No new datasets were generated during the current study.

## Code availability

All statistical analyses were conducted using R (v4.0.3) with the TwoSampleMR (v0.6.30), MendelianRandomization (v0.10.0), MVMR (v0.4.2), and coloc (v5.2.3). The scripts used to perform the analyses and generate the figures are available from the corresponding author upon reasonable request.

## Reporting Guidelines

This study was conducted and reported in accordance with the Strengthening the Reporting of Observational Studies in Epidemiology using Mendelian Randomization (STROBE-MR) guidelines. The completed STROBE-MR checklist is provided as Supplementary Information.

